# Longitudinal 7T MRI reveals volumetric changes in subregions of human medial temporal lobe to sex hormone fluctuations

**DOI:** 10.1101/2022.05.02.490281

**Authors:** Rachel G. Zsido, Angharad N. Williams, Claudia Barth, Bianca Serio, Luisa Kurth, Frauke Beyer, A. Veronica Witte, Arno Villringer, Julia Sacher

## Abstract

The hippocampus and surrounding medial temporal lobe (MTL) are critical for memory processes, with local atrophy linked to memory deficits. Animal work shows that MTL subregions densely express sex hormone receptors and exhibit rapid structural changes synchronized with hormone fluctuations. Such transient effects in humans have thus far not been shown. By combining a dense-sampling protocol, ultra-high field neuroimaging and individually-derived segmentation analysis, we demonstrate how estradiol and progesterone fluctuations affect MTL subregion volumes across the human menstrual cycle. Twenty-seven healthy women (19-34 years) underwent 7T MRI at six timepoints to acquire T1-weighted and T2-weighted images. Linear mixed-effects modeling showed positive associations between estradiol and parahippocampal cortex volume, progesterone and subiculum and perirhinal Area 35 volumes, and an estradiol*progesterone interaction with CA1 volume. We confirmed volumetric changes were not driven by hormone-related water (cerebral spinal fluid) or blood-flow (pulsed arterial spin labeling) changes. These findings suggest that sex hormones alter structural brain plasticity in subregions that are differentially sensitive to hormones. Mapping how endogenous endocrine factors shape adult brain structure has critical implications for women’s health during the reproductive years as well as later in life, such as increased dementia risk following perimenopause, a period of pronounced sex hormone fluctuations.

## Introduction

Ovarian hormones are powerful modulators of neuroplasticity, with animal research offering robust evidence of endocrine regulation of brain morphology on a rapid timescale (1, 2). In the timescale of hours-to-days, rodent and non-human primate studies have demonstrated that estradiol and progesterone elicit modulatory effects on cell proliferation (2), dendritic spine and synapse density (3–6), mitochondrial and synaptic health (7, 8), and myelination (9, 10), suggesting a pivotal role of ovarian hormones in brain structural organization. In humans, the menstrual cycle provides an opportunity to study how endogenous fluctuations in hormones may transiently influence the brain, as estradiol levels increase 8-fold and progesterone levels 80-fold over a period of approximately 25-32 days (11). While a growing number of menstrual cycle studies suggest that ovarian hormone fluctuations do influence brain function and behavior in humans (12–15), it remains less clear how endocrine factors may shape brain structure following the rhythmic nature of the menstrual cycle, and the implications this would have for human adult neuroplasticity.

In this context, the hippocampus is a key region shown to display a remarkable degree of neuroplasticity (16–18) and to be implicated in emotional regulation and cognition (2, 19–21), domains that are susceptible to cycle-dependent fluctuations (22, 23). The hippocampus and extended medial temporal lobe (MTL) are also rich in estradiol and progesterone receptors (24–26), and previous studies suggest that estradiol-dominant menstrual cycle phases are associated with greater hippocampal volume (27–29). Findings have been inconsistent, however, as menstrual cycle studies typically only assess two timepoints and do not directly measure ovarian hormone levels, rather using cycle phase as a proxy for hormone states (28, 29). In a single-subject pilot study (30), we observed that subtle gray matter density changes in the hippocampus paralleled daily fluctuations in endogenous estradiol levels across the menstrual cycle, a dynamic pattern that would have been overlooked with a sparse sampling approach. Thus, the hippocampus and surrounding MTL are promising targets for cycle-related hormonal modulation of structural brain plasticity, but study designs require densely-sampled hormone and neuroimaging data over the timescale of the entire menstrual cycle to best capture intra- and interindividual variability in both cycle variation and brain structure.

Moreover, while most human magnetic reasonance imaging (MRI) studies treat the hippocampus as a homogenous structure, recent advances in neuroimaging allow for more precise delineation of neuroanatomical subregions of the hippocampus and MTL *in vivo* in humans (31–34). This specificity is critical given the unique cytoarchitecture, chemoarchitecture, and circuity of MTL subregions (34–36) that differentially contribute to aging and disease (37, 38). Subregion-specific architecture and circuitry, alongside potential differences in hormone receptor densities (39), suggest that hormone-modulated volumetric changes may manifest differently across the MTL complex. Supporting evidence for the influence of ovarian hormone fluctuations on subregions mainly stems from animal work, with particular emphasis on the cornu ammonis 1 (CA1), a subregion critical for memory integration (40) and in which neuronal loss has been associated with Alzheimer’s disease (41). Estradiol enhances synaptogenesis and spine density in rodent and non-human primate CA1 neurons, while progesterone inhibits this effect (3, 5, 42, 43). Another study in female primates found that estradiol treatment increases pre- and post-synaptic proteins in CA1, while combined estradiol and progesterone decreases these synaptic proteins (44). Another region of key interest is perirhinal Area 35, corresponding to the transentorhinal region and medial perirhinal cortex (31, 45), in which atrophy has been associated with cognitive decline as well as early stages of dementia (45–49). While this subregion has received little attention in regards to endogenous hormone fluctuations, women have greater risk of developing Alzheimer’s disease relative to men, particularly following periods of more pronounced hormone fluctuations later in life, such as after perimenopause (50–52). Thus, understanding how subtle hormone fluctuations may influence CA1 and Area 35 volume in young women would potentially provide insight into underlying mechanisms of risk for cognitive decline in women.

Initial evidence for hormone-modulated changes across human MTL subregions has been observed in a recent single-subject study using 3T MRI, in which the authors observed associations between daily ovarian hormone levels and MTL complex volumes (53). No study has yet to apply a high-density sampling protocol to test whether consistent patterns of hormone-volume associations at the subregion level can be identified in the human hippocampus and surrounding MTL in multiple participants across the menstrual cycle. To shed further light on hormone-associated hippocampal and MTL changes in the female brain, we provide a densly-sampled and detailed ultra-high field neuroimaging dataset in 27 healthy participants, who underwent 7T MRI scanning during six menstrual cycle phases: menstrual, pre-ovulatory, ovulation, post-ovulatory, mid-luteal, and premenstrual. We utilized Automated Segmentation of Hippocampal Subfields software (ASHS (32)), which allows for a sensitive approach to individual differences in MTL subregion morphology (54). Notably, the chosen Magdeburg Young Adult 7T Atlas (31) leverages new information on anatomical variability, resulting in more sophisticated delineation of the boundary between the CA1 and subiculum as well as the parahippocampal cortex, and further segmentation of the perirhinal region into Areas 35 and 36. We developed a systematic protocol for rigorous cycle phase characterization to overcome inaccuracy of menstrual cycle monitoring, a limitation of previous work in this field (55, 56). Based on our pilot study (30), we hypothesized that cycle-related increases in estradiol levels would be associated with increases in whole hippocampus volume. Within the subregions, based on the above-mentioned animal literature, we hypothesized that estradiol levels would be positively associated with perirhinal Area 35 volume, and that there would be an interaction between estradiol and progesterone levels in CA1 volume. The other subregion volumes (CA2, CA3, subiculum, dentate gyrus, Area 36, entorhinal and parahippocampal cortices) were assessed in an exploratory fashion. Given the essential role of the hippocampus and MTL in adult neuroplasticity, these findings may contribute to a better understanding of how endocrine factors shape healthy adult brain dynamics as well as inform more individualized strategies for neuroimaging the MTL complex.

## Results

### Monitoring

All participants were of reproductive age (mean±SD, 25.33±3.64 years) with a healthy body mass index (BMI, 22.37±2.69 kg/m^2^) and regular menstrual cycle length (29.04±2.62 days). Endogenous hormone values, subregion volumes, and whole hippocampus volume (sum of CA1, CA2, CA3, subiculum, dentate gyrus, and remaining tail) were within expected ranges (**Figure 1, Table 1**). For further analyses, hormone values were log-transformed and bilateral subregion volumes were adjusted for total brain volume, and statistical significance was accepted at a Benjamini-Hochberg false detection rate (FDR) corrected threshold of q < 0.05 (see Methods for additional monitoring and statistical details).

**Figure 1:**
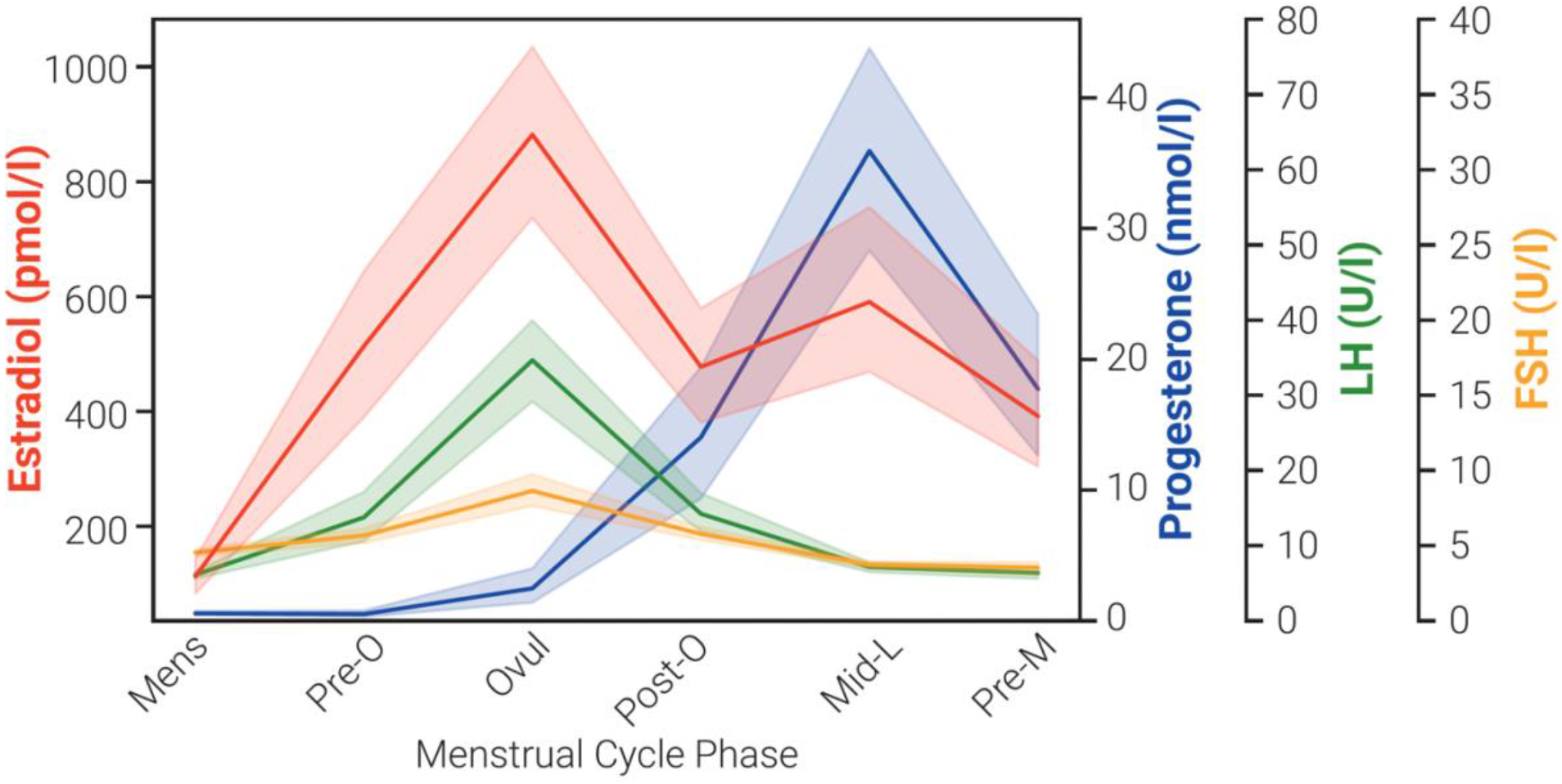
Changes in endogenous levels of estradiol, progesterone, luteinizing hormone (LH), and follicle-stimulating hormone (FSH) across menstrual cycle. Shaded area represents 95% confidence interval for hormone levels at menstrual (Mens), pre-ovulatory (Pre-O), ovulation (Ovul), post-ovulatory (Post-O), mid-luteal (Mid-L), and premenstrual (Pre-M) timepoints.

**Table 1.** Descriptive statistics for each brain region volume, mm^3^. Whole hippocampus is the sum of cornu ammonis [CA] 1, CA2, CA3, subiculum, dentate gyrus, and remaining tail. Standard deviation [SD].

|  | **Mean (mm<sup>3</sup>)** | **SD (mm<sup>3</sup>)** | **Range (mm<sup>3</sup>)** |
| --- | --- | --- | --- |
| **Total Brain Volume** | 1130246.38 | 83996.77 | 412000.00 |
| **Whole Hippocampus** | 5295.36 | 381.71 | 2322.00 |
| **CA1** | 1399.73 | 132.36 | 586.75 |
| **CA2** | 86.49 | 21.22 | 101.25 |
| **CA3** | 257.00 | 37.31 | 167.00 |
| **Dentate Gyrus** | 905.39 | 117.94 | 534.00 |
| **Subiculum** | 2043.41 | 146.34 | 739.25 |
| **Entorhinal Cortex** | 1151.98 | 165.34 | 852.50 |
| **Area 35** | 752.80 | 104.35 | 585.25 |
| **Area 36** | 4046.27 | 543.87 | 1959.25 |
| **Parahippocampal Cortex** | 660.92 | 97.75 | 540.00 |

### Control analyses

Previous work in the field has been critiqued for not taking into account potential hormone-related water shifts or blood flow changes in the brain, which could be misconstrued as hormone-related brain volume changes. While the Magdeburg Young Adult 7T Atlas does exclude the alveus, fimbria, cerebrospinal fluid (CSF), and blood vessels, we additionally assessed CSF and blood flow changes in the hippocampus and did not observe statistically significant associations with estradiol (CSF: β = 4.28, 95% CI = −14 to 6, random effects SD = 22.62, *p* = 0.402) (blood flow: β = 1.23, 95% CI = −3 to 1, random effects SD = 4.19, *p* = 0.195) nor progesterone (CSF: β = 3.67, 95% CI = −1 to 8, random effects SD = 22.43, *p* = 0.134) (blood flow: β = 0.06, 95% CI = −1 to 1, random effects SD = 4.22, *p* = 0.899), giving further confidence that the following results were not erroneously driven by these factors.

### Ovarian hormones and MTL volumes across the menstrual cycle

As hypothesized, linear mixed-effects modeling showed positive associations between estradiol levels and whole hippocampus volume (β = 108.26, 95% CI = 27 to 190, random effects SD = 174.47, *p* = 0.009). In MTL subregions, the addition of the estradiol*progesterone interaction to the model significantly improved model fit only for the CA1 (*X^2^*(1) = 7.691, *p* = 0.006) (**Figure 2A**). Estradiol was positively associated with CA1 volume (β = 42.87, 95% CI = 21 to 65, *p* < 0.001), progesterone was negatively associated with CA1 volume (β = −150.02, 95% CI = −249 to −51, *p* = 0.003), and we observed a significant interaction of estradiol and progesterone with CA1 volume (β = 53.06, 95% CI = 16 to 90, random effects SD = 44.03, *p* = 0.005), such that at higher progesterone levels, the positive effect of estradiol on CA1 volume is attenuated. Progesterone was positively associated with subiculum volume (β = 13.12, 95% CI = 4 to 22, random effects SD = 43.29, *p* = 0.006) (**Figure 2B**) and with Area 35 volume (β = 11.98, 95% CI = 2 to 21, random effects SD = 44.01, *p* = 0.014) (**Figure 2C**). Finally, estradiol was positively associated with parahippocampal cortex volume (β = 24.33, 95% CI = 10 to 39, random effects SD = 32.48, *p* = 0.001) (**Figure 2D**). All four subregions showed significant changes in volume over the six cycle phase timepoints (**Figure 2 column 2**), but such cycle phase effects did not survive correction for multiple comparisons in the whole hippocampus or subiculum. We did not observe any significant relationship between hormones and CA2, CA3, dentate gyrus, entorhinal cortex, or Area 36 volumes (0.196 ≤ *p*’s ≤ 0.884).

**Figure 2:**
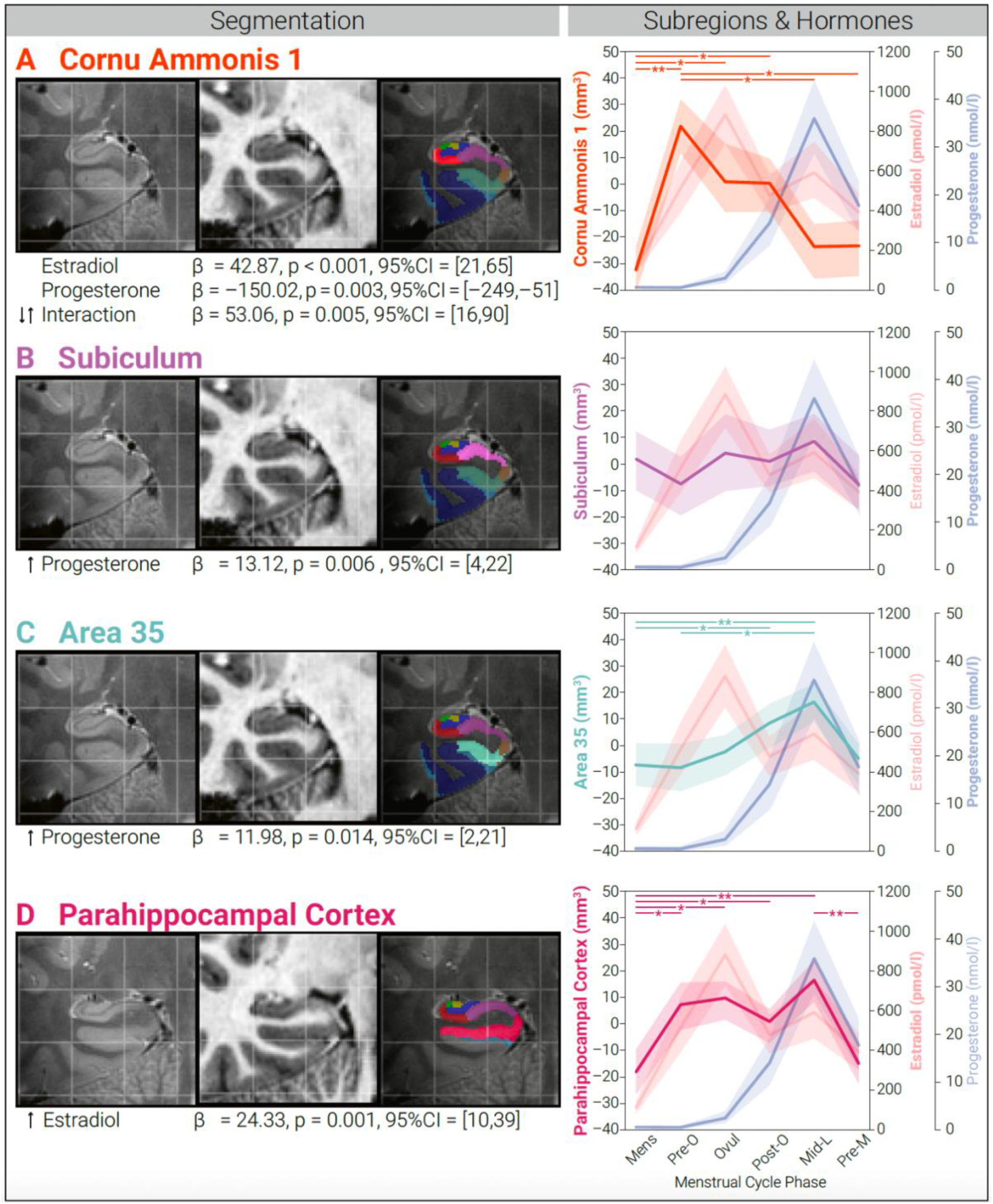
Changes in medial temporal lobe (MTL) volume associated with ovarian hormones across menstrual cycle. *Column 1:* Example T2-weighted image, T1-weighted image, and MTL segmentation for (**A**) cornu ammonis 1, (**B**) subiculum, (**C**) perirhinal Area 35, and (**D**) parahippocampal cortex. *Column 2:* After segmentation, unique associations between ovarian hormones and MTL regions across menstrual cycle. Linear mixed-effects model statistics reported in Column 2. Shaded area represents 95% confidence interval for hormone levels or subregion volumes. Asterisks refer to statistically significant changes in volume over timepoints, \**p*<0.05, \*\**p*<0.005.

## Discussion

In this study, we combined dense hormone sampling with individually-derived high resolution MTL segmentation analysis to demonstrate how estradiol and progesterone fluctuations affect MTL and hippocampal subregion volumes. Parahippocampal cortex, Area 35, subiculum, and CA1 volumes showed significant changes in association with hormone fluctuations across the menstrual cycle. More specifically, estradiol levels were positively associated with parahippocampal cortex volume, and progesterone levels were positively associated with subiculum and Area 35 volume. We also observed an estradiol*progesterone level interaction with CA1 volume. The observed volumetric changes and their unique associations with ovarian hormones, especially progesterone, were concealed when analyzing the hippocampus as a whole, a potential limitation for neuroimaging studies that assess the hippocampus as a single homogenous structure. Finally, we provided further confidence that the observed effects do not erroneously stem from extra-neuronal factors, such as hormone-induced changes in regional blood flow or CSF expansion, as we did not find associations between these factors CSF and ovarian hormone levels. Taken together, our results suggest that ovarian hormones rapidly alter structural brain plasticity in subfields that may be differentially sensitive to hormones.

We were especially interested in CA1 and Area 35 given previously observed selective patterns of neuronal vulnerability to memory impairment in CA1 (41) and Area 35 (45–49), as women are more likely to suffer from cognitive impairment when ovarian hormones rapidly fluctuate, such as during perimenopause (50–52). The hypothesized interaction in CA1 is in line with previous animal work, showing that estradiol enhances synaptogenesis, spine density, and synaptic protein levels in CA1, while subsequent increases in progesterone seem to inhibit this effect (3, 5, 8, 42–44). Moreover, progesterone administration decreases dendritic spines in rodent CA1 neurons, an effect that can be inhibited with a progesterone receptor antagonist (3). Although studies in humans are limited and larger brain volume does not necessarily imply better function, we do know that CA1 plays a distinct functional role in memory integration and inference (40). Clinical studies have also shown that estrogen replacement is associated with maintaining cognitive function in older age while progesterone may counteract the benefit of estradiol’s cognitive enhancing effect (57–63). Thus, our findings are consistent with previous animal and clinical work, suggesting a proliferative effect of estradiol and suppressive effect of progesterone on synaptic plasticity in CA1, a region critical for cognitive processes.

We also observed a positive association between progesterone levels and Area 35 volume, which corresponds to the transentorhinal region and medial perirhinal cortex (31, 45). This region is clinically relevant with regards to aging and disease, with previous work showing that neurodegeneration and atrophy in this region is associated with early stages of dementia and cognitive decline (45–49). Given the increasing focus on the interplay between endogenous hormone fluctuations, the MTL, and the disproportionate risk for Alzheimer’s disease in women (51, 52), we hypothesized that estradiol fluctuations would be associated with Area 35 volumetric changes. The observed progesterone association is a novel finding, as the majority of related literature on hippocampal structural plasticity has thus far focused on estradiol relative to progesterone, and the CA1 and dentate gyrus relative to other MTL regions. Additional emphasis on progesterone in future research is warranted given that levels change approximately 80-fold over the menstrual cycle (11), and that both our current findings and previous work (53) show a complex role of progesterone on MTL subregion volumes. We note that, while the single-subject study (53) also observed both positive and negative associations between progesterone and subregion volumes, our observed progesterone findings occurred in different areas of the MTL, where the other study observed associations with CA2/3, perirhinal, entorhinal, and parahippocampal cortex volumes. These differences may be partially driven by differences in study design such as number of participants, scanner strength, segmentation atlas used, and differences in timepoints, and we therefore encourage replication of the current findings in future studies.

Beyond the hypothesized regions, we also observed positive associations between estradiol and parahippocampal cortex volume and progesterone and subiculum volume. While the effect of menstrual cycle timepoint did not survive correction for multiple comparisons in the subiculum, changes in subiculum and parahippocampal cortex volumes have been observed during pregnancy (64) and after surgical menopause (65), respectively, both of which are times of more extreme changes in ovarian hormones. The subiculum has also been previously shown to display sex-specific volumetric changes in healthy cognitive aging (37) and plays an important role in mediating hippocampal-cortical interactions(66), which are critically involved in cognitive function and emotional regulation, processes influenced by menstrual cycle phase. While we highlight the relevant strengths of the methods used in this manuscript (e.g., atlas with distinction of Area 35, better boundary of subiculum and parahippocampal cortex (31)), future research is required to replicate the current findings in these subregions, as our study clearly demonstrates complex interactions between ovarian hormones and MTL structure as a first step in understanding hormone-induced modulation of brain plasticity at the subregion level.

Although the nature of our MRI study cannot directly evaluate the physiological processes underlying MTL morphology changes, we can speculate on potential hormone-induced molecular and cellular mechanisms that may contribute to mesoscopic changes in regional brain volume. Animal research lends proof that ovarian hormones serve as critical components of cell survival and plasticity, where possible mechanisms at the microanatomical level could include cell proliferation and microglial activation (2, 67), dendritic spine and synapse density (3–6), mitochondrial and synaptic health (7, 8), and myelination (9, 10), which can occur in a manner of hours-to-days. These hormone-induced actions can occur through activation of classical estrogen (ERα and ERβ) and progesterone receptors (PR), of which the MTL complex is dense with (24–26). For example, previous work has shown that ER antagonists and agonists can respectively decrease and increase cell proliferation in the dentate gyrus (68, 69). We note that we and others (44, 53) did not, however, observe volumetric changes in this region across the menstrual cycle, suggesting that the MTL changes we observe may be more driven by synaptic plasticity and remodeling in regions such as the CA1 as opposed to neurogenesis in the dentate gyrus. Indeed, within the hippocampal complex, these three classical receptors appear most prominent in the CA1 (39), where estradiol administration and ER agonists increase synaptic proteins (44, 70), ER antagonists decrease synaptic proteins (71), and progesterone administration and PR antagonists respectively decrease and increase dendritic spines (3, 5). These modulatory effects were not as prominent in the dentate gyrus and other subregions as compared to in CA1 (39, 44, 70, 71). Our study thus encourages future investigation of specific biological microstructural mechanisms underlying observed short-term dynamics of human MTL subregion morphology during times of ovarian hormone fluctuations.

While the rigorous cycle monitoring protocol and individually-derived MTL segmentation analyses serve as strengths of this sufficiently powered longitudinal study, several limitations should be acknowledged. We recruited a healthy population to reflect how endogenous endocrine factors regulate healthy adult brain structural plasticity in the female reproductive years. Therefore, future studies should test whether these hormone-induced changes in MTL subregions are further exacerbated during times of more profound neuroendocrine change, and whether there are behavioral consequences for patient populations. Recent work does suggest that early changes in neuropsychiatric and neurodegenerative disease are better detected in smaller subregions of the MTL rather than with whole structure analysis (46–48, 72), and women are at increased risk for such disorders following ovarian hormone fluctuations, such as depressive disorders (73–77), multiple sclerosis (78–80), dementia (50, 52), and other inflammation-related disorders (81) during the premenstrual phase (73–75, 78, 81), postpartum period (73, 76, 79), and following perimenopause (50, 52, 73, 77, 80). Given the essential role of the MTL in adult synaptic plasticity and the differential contributions of MTL subregions to cognitive functions (40, 82–86), this information may be particularly relevant for plasticity-related disorders such as depression and dementia. Furthermore, while our analyses focused on the hippocampal and MTL complex, ovarian hormones have widespread effects on many brain areas that work in tandem. Subregions have distinct connectivity profiles (35) and future studies should extend the focus to other brain areas to capture a broader network understanding. Given that we saw unique hormone interactions with CA1 and subiculum, and that hippocampal projections to the medial prefrontal cortex (PFC) originate primarily in the CA1 and subiculum (87), we suggest investigation of hormone-modulated changes in structural and functional connectivity between these subregions and the PFC. The PFC also densely expresses hormone receptors (88) and displays hormone-induced structural plasticity (4, 7, 8), with the hippocampal-prefrontal pathway implicated in cognitive and emotional processes (66, 87).

Despite decades of scientific evidence for dynamic interactions between the endocrine and nervous systems, neuroscientific research has largely ignored how endogenous ovarian hormone fluctuations influence human adult brain structural plasticity. This, alongside the current underrepresentation of female samples in neuroscience (55, 89–91), directly limits opportunities for basic scientific discovery and the diversity of human brain health. The MTL region has a remarkable degree of plasticity in response to subtle hormone changes across the female lifespan, and our findings suggest that these changes are detectable at the subregion level. To our knowledge, this is the first study to use ultra-high field neuroimaging to demonstrate how endogenous hormone fluctuations rapidly and transiently alter volume across the MTL complex in multiple participants. We demonstrate the feasibility of such a longitudinal MRI design for creating dynamic and personalized maps of the human brain to inform more individualized strategies for neuroimaging the MTL, which may ultimately lead to better understanding of the manifestation and treatment of hormone-related neuropsychiatric and neurodegenerative diseases.

## Methods

### Participants

Eligible individuals were female, right-handed, 18–35 years old, with a BMI 18.5–29 kg/m2, and without any neurological or psychiatric illness as confirmed with a structured clinical interview. Exclusion criteria were prescription medication or supplement use, tobacco use, positive drug or pregnancy tests, use of hormonal contraceptives, or having been pregnant, postpartum, breastfeeding, or had an abortion within one year of the study. Participants were screened and excluded for DSM-IV Axis I Disorders (92) and Axis II Disorders (93), as well as for presence of premenstrual mood symptoms using the Premenstrual Symptoms Screening Tool (PSST) (94). All participants provided written informed consent after all procedures were fully explained. Forty-one participants were enrolled, of whom two were excluded due to inability to tolerate the 7T MRI scan, eight voluntarily discontinued due to time demands of study, and four were excluded due to irregular cycles, irregularities in bloodwork, or emergency contraceptive pill use after enrollment (included participants N = 27). Of the included participants, twenty completed all six timepoints, two completed five timepoints, one completed three timepoints, one completed two timepoints, and three completed one timepoint (dropout reasons: five due to scheduling conflicts, one used the emergency contraceptive pill, one got an MRI-incompatible retainer); for a total of 138 assessments. The Ethical Committee at the Medical Faculty of Leipzig University approved the study, protocol, and informed consent forms (#077-11-07032011), and the study has been pre-registered at the Open Science Framework (https://osf.io/8mk74/).

### Assessment timing

Participants had a documented history of regular menstrual cycles. The study assessments occurred during six cycle phases: menstrual (<5 days menses onset), pre-ovulatory (≤2 days before ovulation), ovulation (≤24 hours of ovulation), post-ovulatory (≤2 days after ovulation), mid-luteal (6-8 days after ovulation), and premenstrual (≤3 next menses onset). We developed a systematic protocol for rigorous menstrual cycle monitoring and characterization to determine cycle phase timing as follows. After enrollment, participants used an online application (https://www.mynfp.de) to record daily vaginal basal body temperature, menses information, and cycle day and length information. To determine ovulatory timing, participants underwent multiple vaginal ultrasounds to track growing follicle and detect ovulation, completed luteinizing hormone urine tests throughout the ovulatory week, and consulted with a gynecologist. The first assessment phase was randomized across participants, and all other assessments took place in remaining chronological order (menstrual, pre-ovulatory, ovulation, post-ovulatory, mid-luteal, premenstrual).

### Blood Measurements

Serum from fasting-blood samples was collected at every ultrasound and assessment day visit to measure hormones, confirm cycle phase, ensure physical health, and exclude pregnancy or recent drug-intake. Serum was delivered immediately to the hospital laboratory and kept at 5°C until assayed within 24 hours. Estradiol and progesterone concentrations were determined using high performance liquid chromatography-tandem mass spectrometry (LC-MS/MS), and follicle-stimulating hormone and luteinizing hormone concentrations were determined using electrochemiluminescence immunoassay (ECLIA; Roche). Of the 138 assessments, estradiol values for two assessments and progesterone values for one assessment were not included due to pre-analytical error.

### 7T MRI acquisition

Anatomical MRI scans were acquired at the Max Planck Institute for Human Cognitive and Brain Sciences, Leipzig, using a Siemens Magnetom 7T system (Siemens Healthineers, Erlangen, Germany) and 32-channel head array coil (NOVA Medical Inc., Wilmington MA, USA), matched for time of day and without caffeine intake. We acquired high-resolution whole-brain T1-weighted images using an MP2RAGE protocol (repetition time (TR) = 5000 ms; inversion time (TI) 1/2 = 900/2750 ms; echo time (TE) = 2.45 ms; image matrix: 320 × 320 × 240; voxel size 0.7 mm × 0.7 mm × 0.7 mm; flip angle 1/2 = 5°/3°; parallel imaging using GRAPPA with acceleration factor = 2). We acquired T2-weighted imaging slabs perpendicular to the anterior-posterior axis of the hippocampus using a Turbo-Spin Echo Sequence (TR= 16000 ms; TE = 14 ms; image matrix: 384×384; 50 slices; voxel size: 0.5 mm × 0.5 mm × 1 mm; refocusing flip angle = 120°; turbo factor = 8; parallel imaging using GRAPPA with acceleration factor = 2).

### MTL segmentation and volumetry

Background noise removal from uniform T1-weighted MP2RAGE image volumes were done using https://github.com/JosePMarques/MP2RAGE-related-scripts (95). The high resolution T1- and T2-weighted images were then submitted to the Automatic Segmentation of Hippocampal Subfields (ASHS) package (32) using the Magdeburg Young Adult 7T Atlas (31), which has been shown to be more sensitive to individual differences in MTL subregion morphology compared to FreeSurfer (54). ASHS segmentation software uses a fully automated framework at all stages (MRI pre-processing, rigid body transformation alignment of T1- and T2-weighted images, bias correction and refining, etc.), automatically segmenting the MTL in the T2-weighted MRI scans. ASHS documentation, atlases, and software are available at https://sites.google.com/view/ashs-dox/ and https://www.nitrc.org/projects/ashs, with technical details and reliability described further in (32). We performed segmentation and bilateral volume calculations for hippocampal (CA1; CA2; CA3; subiculum; dentate gyrus) and adjacent MTL subregions (entorhinal cortex; parahippocampal cortex; perirhinal cortex [segmented into Area 35 and Area 36]). ITK-SNAP (v3.8) was used for quality assurance (e.g., proper alignment of T1- and T2-weighted images). Quality assurance images were visually assessed by two raters blinded to cycle phase. All code is publicly available at https://github.com/RGZsido/MTLPlasticity2022.

### Total brain volume, CSF, and cerebral blood flow

Total brain volume as well as CSF were calculated using the 7T T1-weighted images and the Segment Data module in CAT12 toolbox of SPM12 MATLAB R2021a, with all overall weighted image quality (IQR) measures > 90%. Cerebral blood flow was calculated using a T1-weighted MPRAGE (TR = 2300 ms, TI = 900 ms, TE = 4.21 ms, flip angle = 9°, FOV = 256 × 256 mm, slices = 176, bandwidth = 240 Hz/px, voxel size = 1×1×1 mm^3^) and a pulsed arterial spin labeling (pASL) sequence (TR = 3000, TI1 = 700 ms, T1S = 1775, TI2 = 1800ms, TE = 13 ms, flip angle = 90°, matrix size = 64 × 64, slices = 24, FOV = 192 × 192 mm, voxel size = 3 × 3 × 4 mm^3^; labeling slab thickness = 100 mm with a gap of 22 mm, 101 pairs of label and control images) (96), acquired on the same assessment days on a 3T Magnetom Verio scanner (Siemens, Erlangen, Germany) using a 32-channel head coil. The pASL data was preprocessed using an inhouse MATLAB analysis pipeline, which included co-registration to the MPRAGE image, motion correction with linear regression, normalization to MNI space, and smoothing with a 2D spatial Gaussian filter of 3-mm FWHM. The final cerebral blood flow values used for the analysis were calculated by pairwise subtraction of labeled and control images by the perfusion model (97).

For the cerebral blood flow analysis, an anatomical region-of-interest (ROI) was created as binary mask of the hippocampus using the WFU PickAtlas toolbox. The mask was resampled to a 3×3×4-mm voxel size to match the pASL images using the coregister:reslice function in SPM12. The preprocessed cerebral blood flow maps were multiplied with the binary mask of the hippocampus and the average cerebral blood flow value (over all voxels within the ROI) was extracted. This was done for all timepoints for each participant.

### Statistical analysis

Assuming a medium effect size (*η2* = 0.06), alpha coefficient of 0.05 and a power of 80 percent, we calculated a total sample size of N = 18 (*a priori* power analysis, G*Power). In the project protocol, we stated that we aimed to include N = 20 healthy participants with all six timepoints. Estradiol and progesterone values were log-transformed prior to analyses. For outlier detection in brain volumes, we flagged bilateral volumes that were three standard deviations from the mean for anatomical inspection by two raters. If upon inspection the volume segmentation maps were unanimously deemed anatomically sound, the volumes were kept to capture reasonable anatomical variation. Of the 138 assessments, we flagged a total of four brain volumes, of which two were determined to be of poor segmentation quality (both for CA1), and were thus removed from further analyses. Bilateral subregion volumes were then adjusted for total brain volume (unstandardized residuals).

For control analyses, we performed linear mixed-effects modeling using the maximum likelihood method of the ‘lmer’ function in the ‘lme4’ R package (v3.5.2 (98)) to assess potential effects of hormones (estradiol and progesterone) on CSF and cerebral blood flow. For main analyses, we used linear mixed-effects models to assess fixed effects of hormones as well as their interaction on whole hippocampus and on each subregion volume. Inclusion of the interaction term was assessed by comparing model fits using the ‘anova’ function. The p-values of the model parameters were calculated via Wald tests and corrected for multiple comparisons using Benjamini–Hochberg procedure (99) controlling for FDR, and were accepted at an FDR-corrected threshold of q < 0.05. For brain volumes that showed significant effects of hormones, we then investigated the fixed effects of cycle phase timepoint modelled as an independent regressor, and performed post-hoc tests using the ‘difflsmeans’ function and the Satterthwaite correction for degrees of freedom. Participants were included as a random factor in all models. The R code for analyses is publicly available at https://github.com/RGZsido/MTLPlasticity2022.

## Acknowledgements

Preparation of this manuscript was supported by a Fellowship from the Joachim Herz Foundation (RGZ), the Branco Weiss Fellowship, Society in Science, National Association for Research on Schizophrenia and Depression (NARSAD) Young Investigator Grant 25032 from the Brain & Behavior Research Foundation (JS), and a Minerva Research Group Grant from the Max Planck Society (JS). The funders of the study had no role in study design, data collection, data interpretation, or writing of the report. The corresponding author had full access to all the data and final responsibility for the decision to submit for publication. We thank Matthias Heinrich for assistance in data acquisition, Toralf Mildner for assistance in pASL preprocessing, and Cornelia Ketscher and Heike Schmidt Duderstedt for assistance with data visualization.

